# Genome report: chromosome-scale genome assembly of the Olive fly *Bactrocera oleae*

**DOI:** 10.1101/2025.07.30.665224

**Authors:** Thorsten E. Hansen, Renee L. Corpuz, Tyler J. Simmonds, Charlotte Aldebron, Charles J. Mason, Scott M. Geib, Sheina B. Sim

## Abstract

The olive fruit fly, *Bactrocera oleae* (Rossi) (Diptera: Tephritidae), is a specialist of fruits of the genus *Olea* and is a major pest of commercial olives due to their adverse impacts to olive production. In support of genomic and physiological research of the olive fly, we sequenced, assembled, and annotated two independent genomes, one from a wild-collected male and one from a wild-collected female. The resulting genomes are highly contiguous, collinear, and complete, attesting to the accuracy and quality of both assemblies. In addition to the autosomes captured as single contigs, the X and Y chromosomes were also captured as evidenced by the X chromosome showing diploid coverage in the female assembly compared to haploid coverage in the male assembly and the Y chromosome being entirely absent from the female assembly. These assemblies represent the first full chromosome-level assembly for *Olive fly*. In addition, a complete genome assembly of a known obligate symbiont to the olive fly, *Candidatus Erwinia dacicola*, was fully captured. The *Ca. E. dacicola* we report here is the most contiguous to date, represented with a gapless chromosome and two separate gapless plasmids. These genome assemblies, along with bacterial symbiont assembly, provide foundational resources for future genetic and genomic research in support of its management as an agricultural pest.

## Introduction

The olive fruit fly, *Bactrocera oleae* (Rossi) (Diptera: Tephritidae), is among the most significant olive pests worldwide. Olive fly is disturbed throughout Mediterranean basin, South and Central Africa, Canary Islands, the Near and Middle East, California, and Central America (Daane & Johnson, 2010). This pest’s range continues to expand alongside olive production with its recent introduction into Hawai’i (Matsunaga et al., 2019). In stark contrast to most other *Bactrocera* pest species (Vargas et al., 2015), olive fly larvae are monophagous feeders of olive fruits in the genus *Olea* (e.g., *O. europaea, O. verrucosa*, and *O. chrysophylla*). Females lay eggs in ripe fruit, and larvae consume the olive mesocarp, spoiling table olives and lowering olive oil quality (Neuenschwander & Michelakis, 1978). Olive fly has been estimated to damage 5% of total olive production, resulting in economic losses of approximately US$ 800 million/year (Bueno & Jones, 2002). Various methods have been used to control and suppress olive fly populations, such as insecticide sprays (e.g., pyrethroids), particle film sprays, pheromone traps, and sanitation of overwintered fruit (Broumas et al., 2002; Margaritopoulos et al., 2008; Wang et al., 2022). However, these controls methods are too costly and time intensive to deploy on broader scales, can select for insecticide resistance, or have off-target effects. Even the highly successful Sterile Insect Technique has been difficult to implement as a part of olive fly pest management due to the challenges of mass-rearing (Johnson et al., 2006).

Aside from the economic importance of olive fly, the specialist nature of this species is peculiar within the *Bactrocera* genus. Olive fly’s unique host plant use has resulted in a partnership with an unculturable, extracellular, obligate gut symbiont, *Candidatus Erwinia dacicola* (Ben-Yosef et al., 2010; Capuzzo et al., 2005). *Ca. E. dacicola* allows olive flies to feed on unripe olives (Ben-Yosef et al., 2015), provides amino acids or converts inaccessible nitrogen for the host (Ben □ Yosef et al., 2014), and its presence leads to higher attempts of oviposition compared to antibiotic treated flies (Jose et al., 2019). Other tephritid species do not possess as rigid a gut symbiosis, suggesting a more stringent degree of specialization and partnership between species. These observations of the olive fly and its symbiont imply a need for greater genomics resources to help develop and improve present management strategies and provide insight into the evolution of the host-microbial partnership.

In support of foundational and applied research of the olive fly, we sequenced and assembled two chromosome-level genomes, one male and one female. Contig assemblies were generated using PacBio HiFi data from a single wildtype male and a single wildtype female. Enriched chromosome conformation contact data was generated from a second male and female which validated that seven of the contigs of the male assembly and four of the contigs of the female assembly were at a chromosome scale. Nucleotide alignments of the two genomes revealed high-collinearity and parity between the two assemblies and the two sex chromosomes, X and Y. Estimations of genome completeness using BUSCO analysis of the genome and structural annotations revealed both genomes and gene sets achieved greater than 99% of the expected genes in the Brachycera protein set (odb12). Additionally, a complete and continuous genome of the obligate symbiont *Ca. E. dacicola* was produced in tandem. Our methods provide a framework for producing platinum-quality, contiguous, chromosome-scale genome assemblies from single insects with their residing symbionts.

## Methods and materials

### Source material

Infested olive (*Oleae europaea*) fruits were collected at the University of Hawai’i Lalamilo Research Station in Hawai’i County (<u>20.019814 N, 155.677155 W</u>) and returned to the United States Department of Agriculture Agricultural Research Service Pacific Basin Agricultural Research Center in Hilo, Hawai’i, USA. Olive fruits were held to allow larval development and pupation, and adults were maintained in the laboratory with separate water and food source in a 3:1 mixture granulated sugar and yeast hydrolysate. Two olive fly adult males and two adult females were obtained as pairs from this collection and flash frozen in liquid nitrogen for genomic library preparation. Additional olive flies were collected taxonomic identification vouchers. Both a female (Figure 1a) and a male (Figure 1b) were collected immediately after eclosion (∼ 24 hrs), cold shocked, and photographed for the taxonomic identification vouchers.

**Figure 1:**
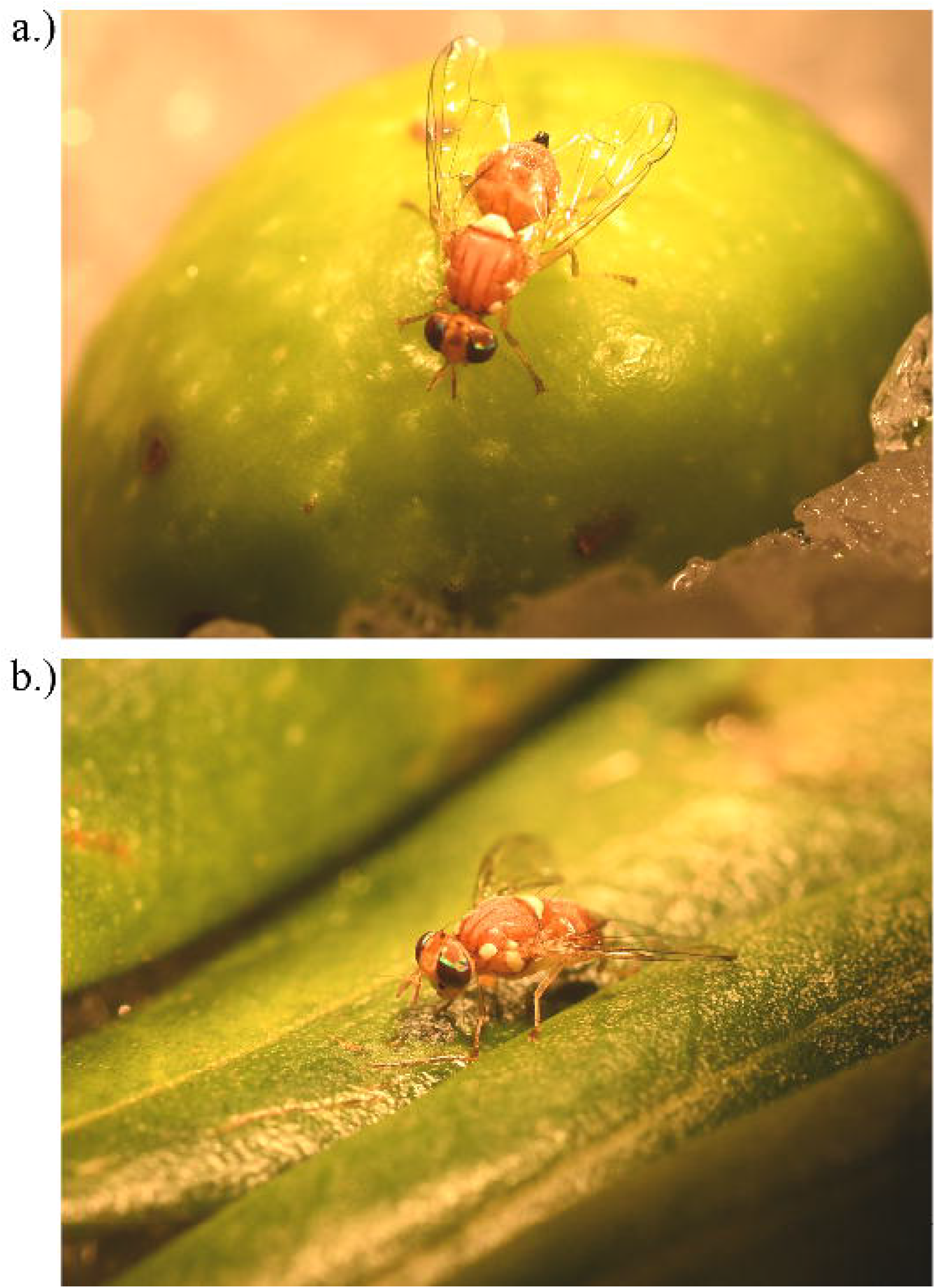
Photo vouchers of adult olive flies. a) This photo of a female adult olive fly was the offspring of wild olive flies collected from oviposited olive fruits. b) This photo of a male adult olive fly was the offspring of wild olive flies collected from oviposited olive fruits.

### Library preparation and sequencing

The whole body of one olive fly male and one female were separately homogenized into a fine powder while kept frozen using a SPEX SamplePrep 2010 Geno/Grinder (Cole Parmer, Metuchen, New Jersey, USA) and underwent HMW DNA extraction using the fresh or frozen tissue protocol of the Qiagen MagAttract HMW DNA Kit (Qiagen, Hilden, Germany). Following isolation, the HMW DNA was sheared to a mean fragment length of 20 Kb with a Megaruptor 3 (Diagenode, Dennville, New Jersey, USA) and prepared into a PacBio SMRTBell library using the SMRTBell Express Template Prep Kit. 3.0 (Pacific Biosciences, Menlo Park, California, USA). After DNA isolation, shearing, and library preparation, samples were purified using solid-phase reversible immobilization beads (SPRI beads) (DeAngelis et al., 1995) and quantified using fluorometry and spectrophotometric absorbance ratios (DeNovix Inc., Wilmington, Delaware, USA). Fragment length distributions after each step were determined by Femto Pulse or Fragment Analyzer (Agilent Technologies, Santa Clara, California, USA). The resulting libraries were sequenced separately on a PacBio Sequel IIe system using a 30-hour movie length on a 8M SMRTCell. Raw subreads were converted to HiFi data using the PacBio SMRTLink software v.10.1.

Concurrent to HiFi sequencing, the second adult, for both male and female, was used to prepare an enriched chromosome conformation capture (HiC) library. Tissue from whole body of one adult olive fly was homogenized in 1× PBS and nuclei were crosslinked in a 16% formaldehyde solution. Following crosslinking, the sample was lysed and digested using the restriction enzymes DdeI and DpnII. To enrich the sample for proximity ligated fragments, a biotin-labeled dATP fill-in step was performed prior to proximity ligation so fragments can be recaptured and enriched downstream. Following proximity ligation, a crosslink reversal step was performed followed by two DNA purification steps using SPRI beads, the removal of biotin from unligated ends, and another DNA purification step. The sample was then sheared using a Bioruptor Pico (Diagenode) targeting 400-450 bp. After shearing, the sample was size-selected using SPRI beads, biotinylated ligation products were recaptured, and the sample was prepared into a short-read sequencing library using the NEBNext Ultra II DNA Library Prep Kit (New England Biolabs, Ipswich, Massachusetts, USA) and Element Adept Kit (Element Biosciences, San Diego, California, USA). The final libraries were sequenced on a partial 2×150 flow cell on the Element AVITI System (Element Biosciences). Following sequencing, raw reads were base called using bases2fastq v.1.3.0.

### Olive fly genome assembly, assessment, contaminant removal and annotation

Prior to assembly, adapter-contaminated HiFi reads were removed from the both the male and the female HiFi read pools using FCS-Adaptor and HiFiAdapterFilt (Sim et al., 2022) (https://github.com/ncbi/fcs). The resulting HiFi reads were used to separately assemble the male and the female data into contigs using HiFiASM (male: v 0.19.3-r572, female: 0.24.0-r702) (Cheng et al., 2021; Cheng et al., 2022). The contig assemblies was subsequently purged of duplicate contigs using PurgeDups (Guan et al., 2020), and the duplicate purged contig assemblies served as the respective references to map male and female HiC reads using BWA-mem 2 (v2.2.1) (Vasimuddin et al, 2019). The resulting mapped files were filtered for artifact PCR duplicates using Picard (v.3.2.0) (Picard2019toolkit, 2019 https://github.com/broadinstitute/picard). Contact maps were generated from the de-duplicated mapped reads using the YaHS pipeline (Zhou et al., 2023). Visualization of the contact maps and minor manual editing was achieved using Juicebox (v.2.15) (Durand et al., 2016). Minimap2 (v.2.22-r1101) was used to map HiFi reads back to the contig assembly and calculate coverage of each contig, the –auto function and –genome mode of BUSCO (v.5.2.2) was used to select the appropriate taxon database and estimate genome completeness, and BLAST+ and Diamond were used to perform nucleotide alignments to the NCBI nucleotide database (accessed February 2022) and UniProt protein database (accessed March 2020) respectively (Buchfink et al., 2021; Camacho et al., 2009; Li, 2018; Manni, Berkeley, Seppey, Simão, et al., 2021; Manni, Berkeley, Seppey, & Zdobnov, 2021). The resulting outputs of minimap2, BUSCO, BLAST+, and Diamond were summarized and visualized using Blobtools2 v and blobblurb (Challis et al., 2020); (https://github.com/sheinasim/blobblurb). Additional taxonomic assignments of contigs and contig fragments was performed using the FCS-GX(Astashyn et al., 2024). The confluence of the taxonomic assignments based on nucleotide alignment, protein alignment, and FCS-GX was used to identify contigs assigned to *Ca. E. dacicola* and putative bacterial plasmids and remove non-Arthropod contigs from the assembly. The male assembly was submitted to the National Center for Biotechnology Information (NCBI) and currently represents the RefSeq genome for this species. Genome annotations for the male genome was performed using the NCBI Eukaryotic Genome Annotation Pipeline (EGAP) (Thibaud-Nissen et al., 2013). The female assembly was also submitted to NCBI and was annotated by the NCBI EGAPx pipeline.

### Olive fly mitochondrial genome

All potential copies of the mitochondrial genome in both assemblies were identified using the MitoHiFi pipeline (Uliano-Silva et al., 2023). MitoHiFi implemented a BLAST search for contigs that have a high similarity to another publicly available mitochondrial genome of olive fly, (NCBI RefSeq accession NC_005333.1) (Camacho et al., 2009), selecting the contig with the greatest similarity and checking for circularization. Mitochondrial genes were then structurally annotated using intervals from the same mitochondrial genome used in the BLAST search through the MitFi annotation program in the MitoFinder pipeline (Allio et al., 2020; Jühling et al., 2012). A representative mitochondrial genome for each assembly was submitted to NCBI with the nuclear genomes.

### *Candidatus Erwinia dacicola* genome assessment and annotation

Three contigs assigned to *Ca. E. dacicola* were isolated from the male assembly and the identity was verified using BLASTN within the BLAST+ program (v 2.13.0) (Camacho et al., 2009). Each individual contig fasta was indexed and assembled reads were aligned back with BWA-MEM (v 0.7.17) (Li, 2013). The bam files were then coordinate sorted, indexed, and used to calculate coverage and mean depth for each contig using SAMtools (v 1.17) (Danecek et al., 2021). Total assembly quality assessment metrics for the *Ca. E. dacicola* were produced with QUAST (v 5.2.0) (Gurevich et al., 2013). Circlator (v1.5.5-docker5) was used to detect and trim the overlap to circularize the assembly from the corrected and trimmed reads and the assembly file (Hunt et al., 2015). Ori-Finder (v 2022) was used to detect the origin of replication in the assembly (Dong et al., 2022).

The *Ca. E. dacicola* contig assembly was checked for gaps with bioawk (https://github.com/lh3/bioawk). Further quality assessment and estimation of completeness of the *Ca. E. dacicola* genome was done with CheckM (Parks et al., 2015) and BUSCO (Manni, Berkeley, Seppey, Simão, et al., 2021; Manni, Berkeley, Seppey, & Zdobnov, 2021). BUSCO estimates the completeness and redundancy of genomes based on universal single-copy orthologs. BUSCO (v 5.4.5), using the –auto-lineage-prok flag, produced completeness estimation summaries for the generic domain of Bacteria and the more specific order of Enterobacterales. CheckM evaluates completeness, contamination, and strain heterogeneity of bacterial genomes with a limited number of marker genes conserved across the domain of Bacteria. We used the lineage-specific workflow in CheckM (v 1.2.2) to produce the statistics. The NCBI Prokaryotic Genome Annotation Pipeline (PGAP) (v 2023-10-03.build7061) (Tatusova et al., 2016) was used to annotate the *Ca. E. dacicola* genome.

## Results and discussion

### Olive fly male and female genome assembly metrics

PacBio HiFi sequencing yielded extremely high-quality genomes for both male and female olive fly. For the male, 2,640,135 raw reads (37.6 Gb of raw data) were obtained, of which 2,640,088 (99.99% of reads) were retained after filtering. For the female, 2,524,582 raw reads (41.0 Gb of raw data) were obtained, of which 2,524,553 (99.99% of reads) were retained after filtering. K-mer analysis of the final assemblies relative to the PacBio HiFi reads used to create the contig assemblies reported a raw quality value (QV) and adjust QV score of 70.022 and 68.778 for the male, and 62.713 and 60.557 for the female, respectively. This demonstrates that the final genomes have high consensus accuracy relative to the sequence data for both male and female assemblies.

The male olive fly genome was sequenced to 88× coverage and resulted in a final assembly size of 468.783 Mb (Figure 2a, Table 1). The final assembly is comprised of seven contigs representing five autosomes, an X and a Y chromosome, a mitochondrial genome, and twenty-four unplaced contigs. Each chromosome is represented by a single contig that was validated by HiC contacts (Figure 2b) and the unplaced contigs represents 5.561 mb (.001% of total assembly). Analysis of completeness of the genome using Brachycera BUSCOs revealed 98.6% of expected genes found in single copy, 0.37% duplicated, 0.44% fragmented, and 0.98% missing out of the expected 5,912 Brachycera orthologs (Figure 2a, Table 2). BlobTools analysis showed no signs of contaminants in the final assembly (Figure 2c). Genome annotation of the male olive fly assembly using the NCBI EGAP detected a total of 12,391 protein-coding genes, 3,571 noncoding genes, 986 non-transcribed pseudogenes, 4,773 genes with variants, and no transcribed pseudogenes, immunoglobulin/T-cell receptor gene segments, or other genes outside of the listed categories (Table S1).

**Figure 2:**
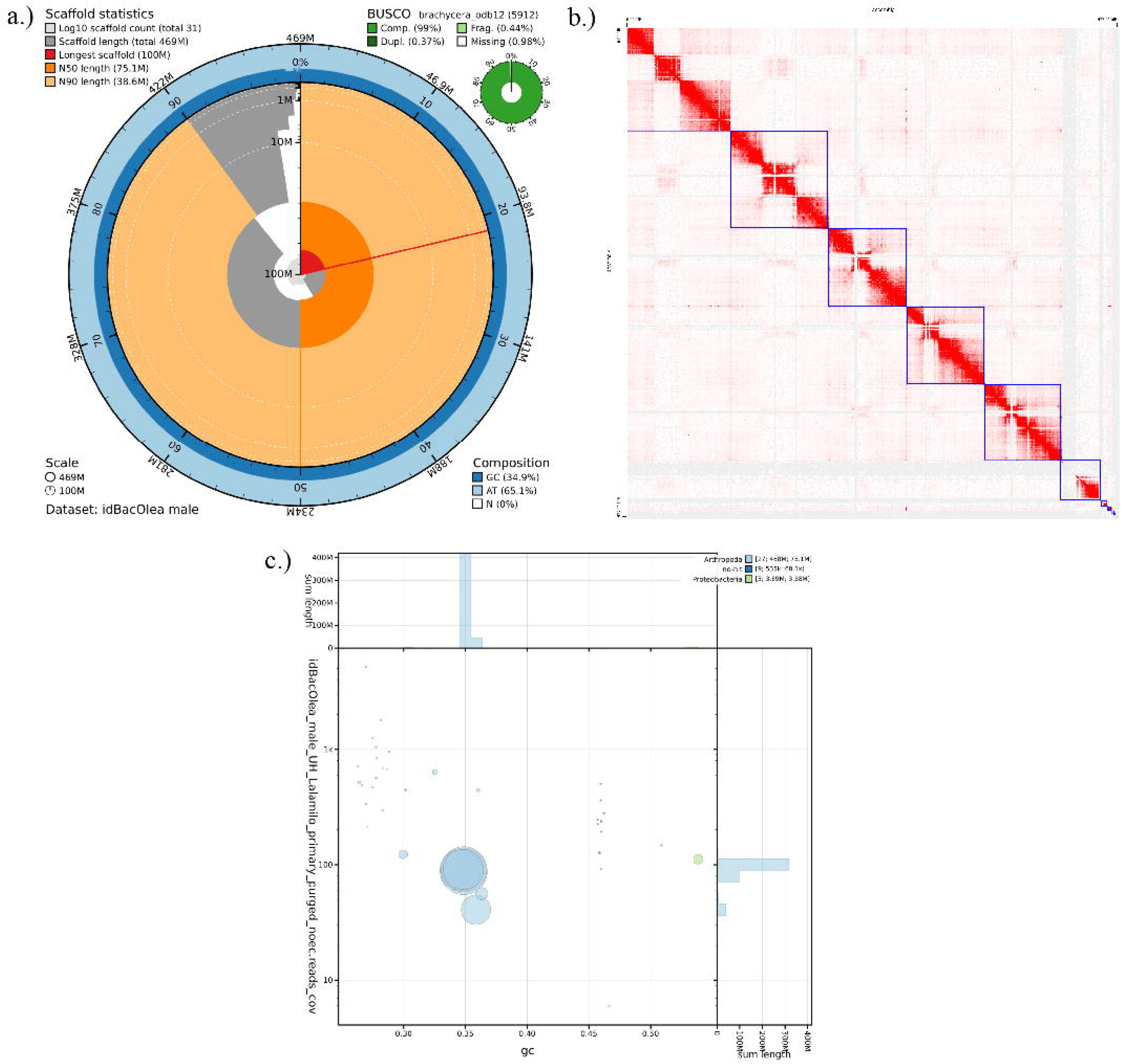
Statistics of the olive fly male genome assembly. a) Snail plot visualization of the contig assembly with BUSCO assessment using Brachycera conserved orthologs. b) Hi-C contact map indicates that contigs can be grouped in seven major scaffolds (blue squares). c) Coverage and GC content of taxon-annotated contigs showing presence of *Candidatus Erwinia dacicola* (green).

**Table 1:** Assembly statistics for the olive fly male and female genome assemblies in comparison to previously *published* olive fly assemblies.

| Species | Genome size (Mb) | Total ungapped length (Mb) | Number of contigs | Scaffold N50 (Mb) | L50 | No. scaffolds | Assembly level | GC content | Reference |
| --- | --- | --- | --- | --- | --- | --- | --- | --- | --- |
| Olive fly | 468.7 | 468.7 | 31 | 75.1 | 3 | 31 | Chromosome | 34.9 | Present study - male |
| Olive fly | 455.396 | 455.394 | 58 | 74.3 | 3 | 47 | Chromosome | 34.9 | Present study - female |
| Olive fly | 484.9 | 458.4 | 48,617 | 4.6 MB | 31 | 38,160 | Scaffold | 34.5 | Bayega et al. 2020 - pooled male & female |
| Olive fly | 334.4 | 330.6 | 100,730 | 10.5 kb | 6,242 | 68,736 | Scaffold | 34.5 | Vicoso & Bachtrog 2015 - pooled male & female |

**Table 2:** BUSCO analysis for the olive fly genome male and female assemblies in comparison to other available genome assemblies.

| Complete [single, duplicated] | Fragmented | Missing | Lineage | Gene set size | Reference |
| --- | --- | --- | --- | --- | --- |
| 98.58 [98.21, 0.37] | 0.44 | 0.98 | brachycera_odb12 | 5912 | Present study - male |
| 98.5 [98.19, 0.34] | 0.46 | 1.01 | brachycera_odb12 | 5912 | Present study - female |
| 98.8 [98.1, 0.7] | 0.7 | 0.5 | brachycera_odb12 | 5912 | Bayega et al. 2020 |
| 73.6 [73.5, 0.1] | 12.3 | 14.2 | brachycera_odb12 | 5912 | Vicoso & Bachtrog 2015 |

**Table 3:**
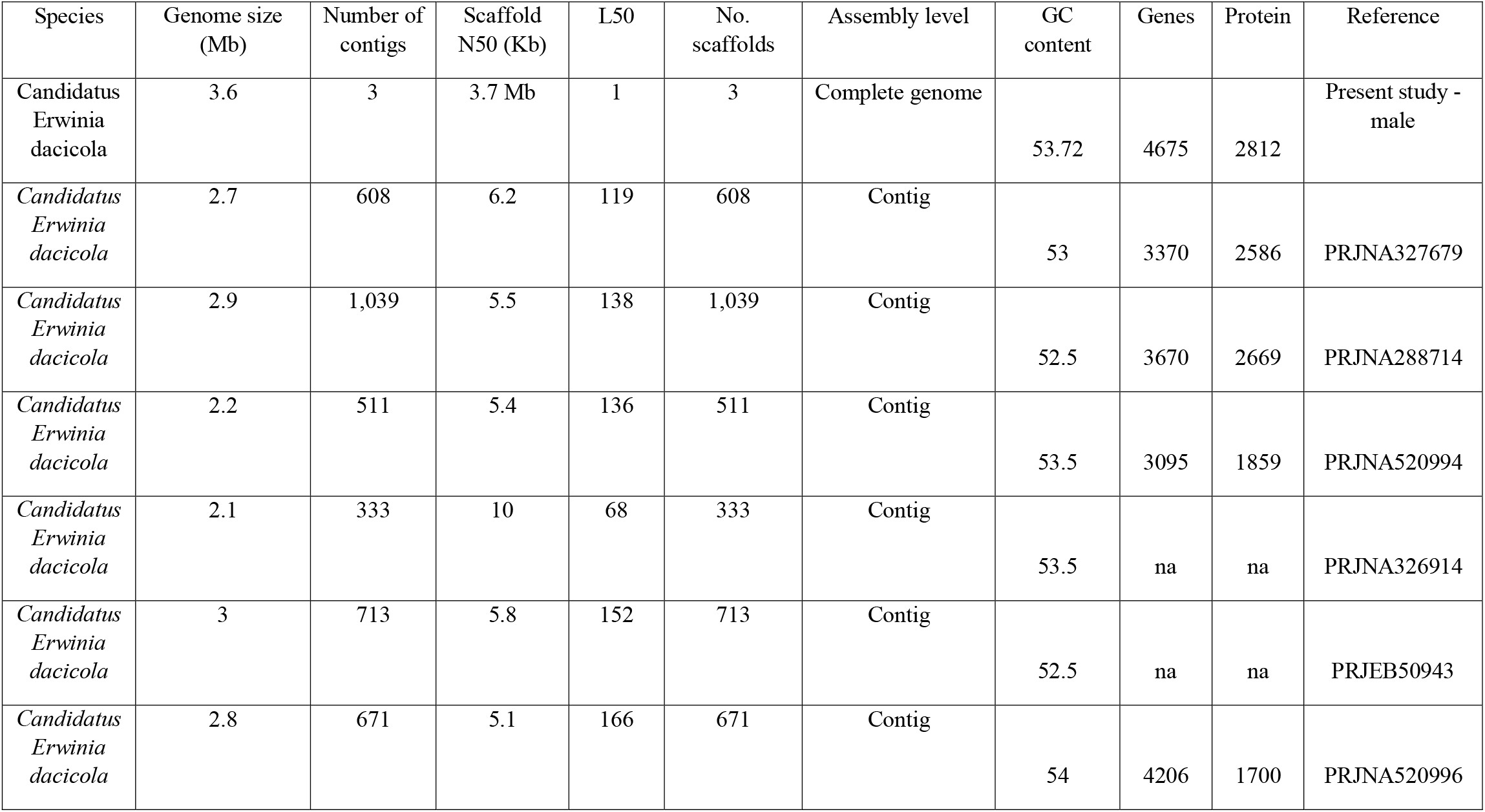
Assembly statistics for the *Candidatus Erwinia dacicola* genome assembly in comparison to other public *Ca. E. dacicola* genome assemblies.

| Species | Genome size (Mb) | Number of contigs | Scaffold N50 (Kb) | L50 | No. scaffolds | Assembly level | GC content | Genes | Protein | Reference |
| --- | --- | --- | --- | --- | --- | --- | --- | --- | --- | --- |
| Candidatus *Erwinia dacicola* | 3.6 | 3 | 3.7 Mb | 1 | 3 | Complete genome | 53.72 | 4675 | 2812 | Present study - male |
| *Candidatus Erwinia dacicola* | 2.7 | 608 | 6.2 | 119 | 608 | Contig | 53 | 3370 | 2586 | PRJNA327679 |
| *Candidatus Erwinia dacicola* | 2.9 | 1,039 | 5.5 | 138 | 1,039 | Contig | 52.5 | 3670 | 2669 | PRJNA288714 |
| *Candidatus Erwinia dacicola* | 2.2 | 511 | 5.4 | 136 | 511 | Contig | 53.5 | 3095 | 1859 | PRJNA520994 |
| *Candidatus Erwinia dacicola* | 2.1 | 333 | 10 | 68 | 333 | Contig | 53.5 | na | na | PRJNA326914 |
| *Candidatus Erwinia dacicola* | 3 | 713 | 5.8 | 152 | 713 | Contig | 52.5 | na | na | PRJEB50943 |
| *Candidatus Erwinia dacicola* | 2.8 | 671 | 5.1 | 166 | 671 | Contig | 54 | 4206 | 1700 | PRJNA520996 |

The female olive fly genome was sequenced to 91× coverage and a total assembly size of 455.396 Mb (Figure 3a, Table 1). The female reference genome is represented by 5 autosomes, an X, a mitochondrial genome, and 40 unplaced contigs. Chromosomes 4 and 5 are each represented by a single contig, chromosomes 2, 3, and 6 are represented by one scaffold made of two contigs each, and the X chromosome was the most fragmented (Figure 3b). Analysis of genome completeness using Brachycera BUSCOs revealed 98.19% of expected genes found in single copy, 0.34% duplicated, 0.46% fragmented, and 1.01% missing out of the expected 5,912 Brachycera orthologs (Figure 3a, Table 2). As with the male assembly, the only non-Arthropod contigs belonged to the *Ca. Erwinia dacicola* associated contigs which were removed from both assemblies prior to NCBI submission (Figure 3c). For the female assembly, structural annotation identified 12,975 protein-coding genes, 2,960 non-coding genes, 295 non-transcribed pseudogenes, 5,404 genes with variants, and no transcribed pseudogenes, immunoglobulin/T-cell receptor gene segments, or other genes outside of the listed categories (Table S1).

**Figure 3:**
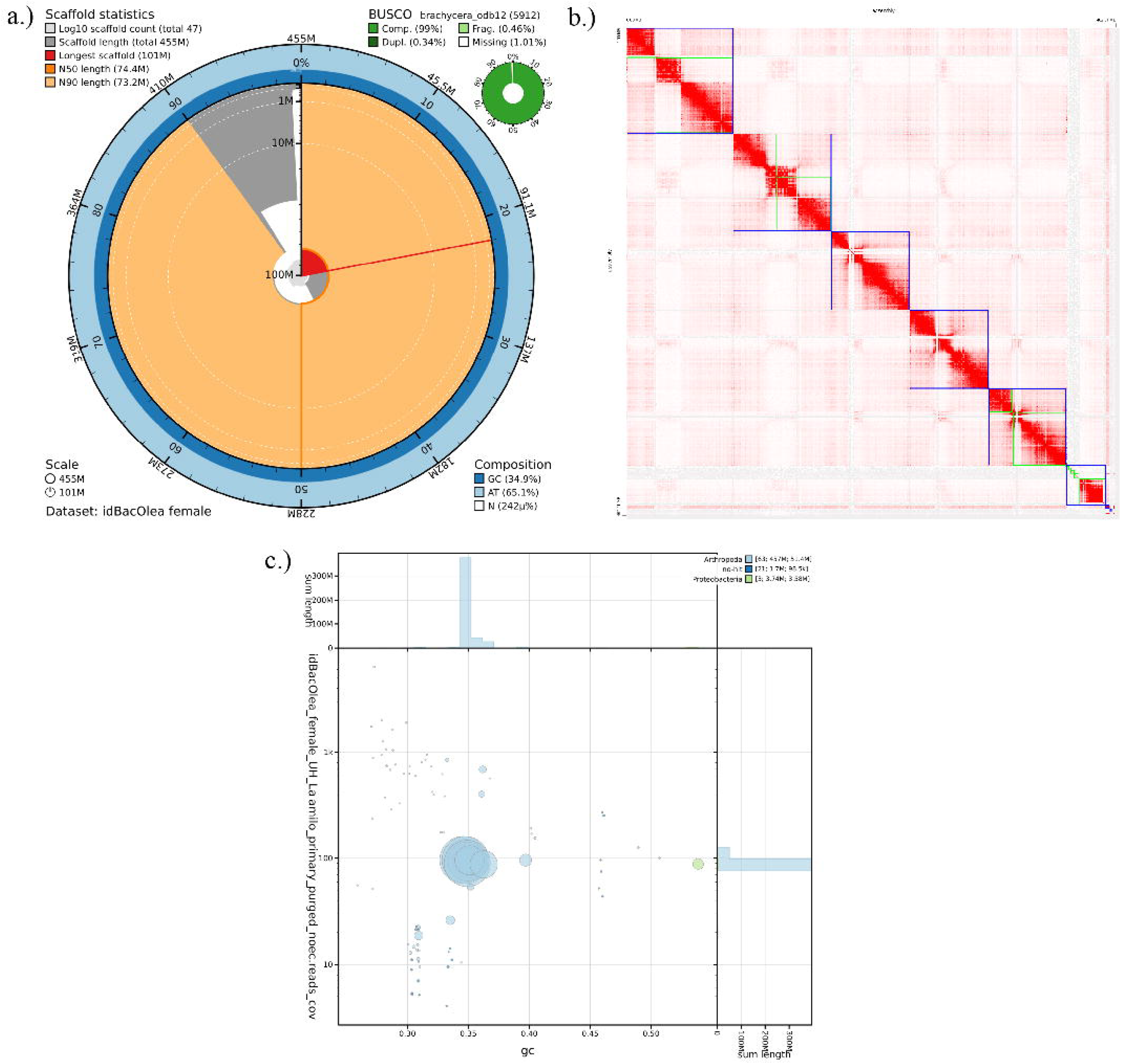
Statistics of the olive fly female genome assembly. a) Snail plot visualization of the scaffold assembly with BUSCO assessment using Brachycera conserved orthologs. b) Hi-C contact map indicates that contigs (green squares) can be grouped in six major scaffolds (blue squares). c) Coverage and GC content of taxon-annotated contigs showing presence of *Candidatus Erwinia dacicola* (green).

The male and female assembles generated in our study are comparable to the two other published olive fly genomes (Bayega et al., 2020; Vicoso & Bachtrog, 2015), but are improved on some important metrics. The genomes we report are the same size and ungapped length as the previously published genomes (Table 1), but our assemblies our comprised of significantly fewer contigs and are assembled to a chromosomal context. Of the previously published olive fly genomes, one was not annotated so statistics concerning completeness and coding sequences in the genome are unavailable (Vicoso & Bachtrog 2015). In comparison the genome produced by Bayega et al. (2020), the assemblies we produced have increased completeness using BUSCOs from the suborder Brachycera.

### Comparison of male and female assemblies and sex chromosome determination

Alignment between the chromosomes of the male and female olive fly assemblies showed high collinearity and congruency between both genomes (Figure 4). The overall difference between the male and the female assembly is miniscule (< 2% difference in assembly size between the female assembly and the male assembly without the Y chromosome) (Table 1). Detection of complete BUSCO orthologs from both the genome (Table 2) and annotated gene set (Table S1) provides support for a high-level of completeness in terms of gene space.

**Figure 4:**
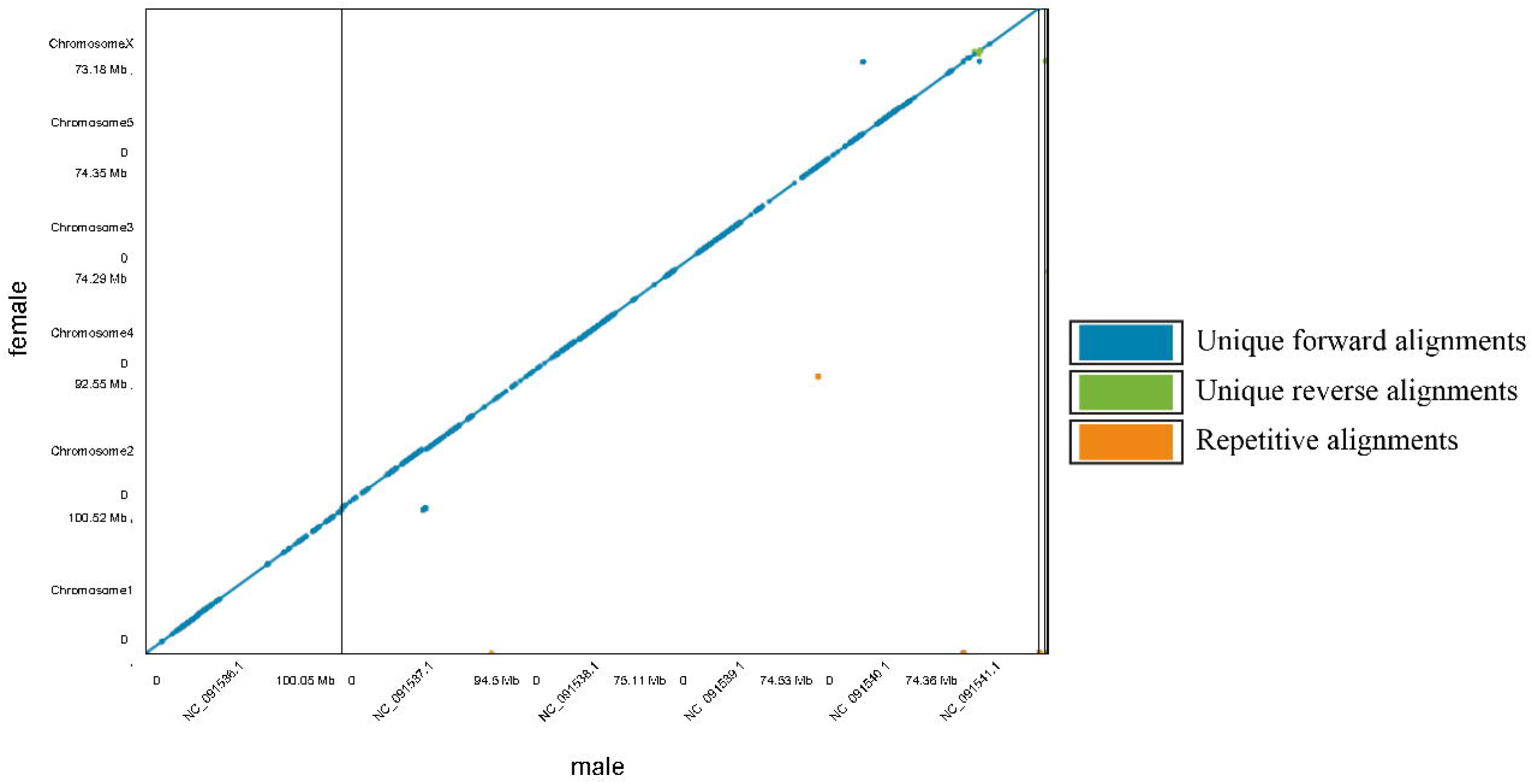
Whole-genome nucleotide alignment between the final male and female olive fly genome assemblies.

The alignment also revealed that the X chromosome displayed high collinearity like the autosomes. The chromosome in the male assembly that did not have a counterpart in the female assembly was assigned as the Y chromosome. From the assemblies the half coverage of the X chromosome in the male (while being similar coverage to the autosomes in the female) along with absence of the Y chromosome in the female support the sequencing and assembly of the sex chromosomes (Table S2).

### Olive fly symbiont *Candidatus Erwinia dacicola* genome assembly metrics

*Ca. E. dacicola*’s genome was comprised of one complete major chromosomal contig (3.603,634 Mb) and two smaller plasmid contigs. Coverage was >99.99% and mean depth was 100.88×. There were zero gaps across the genome assembly. The genome had a GC content of 53.72%. In consideration that this was likely a complete bacterial genome, we checked for circularization with Circlator. Using the HiFi reads from the male olive fly genome assembly, both putative plasmids were circularized, but the program was unable to circularize the *Ca. E. dacicola* major contig. However, we were able to identify an origin of replication (oriC) using Ori-finder located between 1733100 – 1733565 bp at a size 466 bp. OriC’s distance from dnaA (protein that initiates replication), which is closer to the start of the genome (346879 – 346876 bp), combined with its presence near the approximate middle of the genome suggest that *Ca. E. dacicola*’s genome is linear. This is uncommon among bacteria, but not unprecedented (Ochman, 2002; Volff & Altenbuchner, 2000). The *Ca. E. dacicola* assembly also includes plasmids matching *Tatumella* sp. TA1 plasmid pTA1(71,221 bp) and *Citrobacter braakii* strain F217 plasmid pF217-5 (43,375 bp). The first plasmid matched the *Tatumella* sp. TA1 plasmid pTA1 with 100% query coverage and an identity of 99.95%, coverage >99.99%, and mean depth was 86.47×. The second plasmid matched the *Citrobacter braakii* strain F217 plasmid pF217-5 with 92% query coverage and an identity of 96.62%. The second plasmid had coverage 99.99% and mean depth was 7.17×.

Quality assessments indicate a high-quality and mostly complete genome and generic bacterial domain assessment from BUSCO indicated a near complete genome. There were 123 single copy BUSCOs (99.2%) and only one missing (0.8%). The BUSCO analysis performed at the order of Enterobacterales also indicated a complete genome: 423 single copy BUSCOs (96.1), three duplicated BUSCOs (0.7%), two fragmented BUSCOs (0.5%), and twelve missing BUSCOs (2.7%). Specific missing BUSCOs listed in Table S3. CheckM reported a marker gene completeness of 82.52%, with 1.39% contamination, and 87.5% strain heterogeneity. The high strain heterogeneity suggests nucleotide differences within the *Ca. E. dacicola* population we sequenced from.

In total, for the *Ca. E. dacicola* assembly, 4,675 genes were annotated. Approximately 80 coded for RNAs: 55 tRNAs, 20 rRNAs (six – 5S, seven – 16S, seven – 23S), and five ncRNAs. In terms of coding sequences, only 2,812 are protein-coding.

Overall, in comparison to previously published *Ca. E. dacicola* genomes (Table S4) this genome is the most contiguous. Of the assemblies, it is the only one whose chromosome was fully assembled in a singular contig and contains a putatively complete associated plasmid. It encodes for more genes and has more identifiable genomic features (e.g., pseudogenes) than the previously published genomes (BioProject accession numbers: PRJNA327679, PRJNA288714, PRJNA520994, PRJNA326914, PRJNA520996, PRJEB50943).

## Conclusions

The reported olive fly reference genomes are highly contiguous (male N50 = 75.1Mb; female N50 = 74.4 MB) and complete (male complete BUSCOs = 98.6%; female complete BUSCOs 98.5%) chromosome-scale assemblies which will support the development of future applied pest management and as well as foundational research. This high-quality genome assembly contains an identified X (∼ 37.23 Mb) chromosome and Y chromosome (∼ 6 Mb) which can facilitate sex-specific pest management strategies. These host reference genomes can also help inform the processes underlying host plant adaptation and utilization alongside identifying genes that can be for fruit fly management. Furthermore, due to the olive fly’s intertwined relationship with their obligate symbiont (Ben-Yosef et al., 2010; Ben-Yosef et al., 2015; Ben□Yosef et al., 2014), the high-quality references obtained of both the insect host and bacteria can support future analyses investigating the maintenance and evolution of these interactions. This study represents the first public olive fly genome to be assembled at chromosome level with sex chromosomes and will provide valuable information to support pest management programs.

## Data availability

The olive fly genome assemblies are NCBI BioProject Accessions: PRJNA1089778 and PRJNA1275571, male and female, respectively. GenBank accession is GCA_042242935.1 (male) and GCA_051013405.1 (female). RefSeq accession is GCF_042242935.1 (male). PacBio HiFi reads are available on the NCBI Sequence Read Archive (SRA) at accession SRX23996476 (male) (ToLID: idBacOlea1) and SRR33945991 (female) (ToLID: idBacOlea3). Illumina sequences used for HiC scaffolding are available on SRA at SRX23996477 (male)(ToLID: idBacOlea2) and SRR33945990 (female) (ToLID: idBacOlea4). The *Ca. E. dacicola* genome assemblies are NCBI Bioproject ID <u>PRJNA1090968</u>.

## Acknowledgments

This material was made possible, in part, by a Cooperative Agreement from the APHIS. It may not necessarily express APHIS’ views. USDA is an equal opportunity employer. Mention of trade names or commercial products in this publication is solely for the purpose of providing specific information and does not imply recommendation or endorsement by the USDA.

## Funding

This study was funded by the USDA Agricultural Research Service (ARS) with project “Advancing Molecular Pest Management, Diagnostics, and Eradication of Fruit Flies and Invasive Species” (2040-22430-028-000-D) as well “Integrative identification methods for Bactrocera fruit flies” (3.0624). This research used resources provided by the SCINet project and/or the AI Center of Excellence of the USDA Agricultural Research Service, ARS project numbers 0201-88888-003-000D and 0201-88888-002-000D.

## Supplementary data

Table S1: Annotation statistics from the NCBI eukaryotic genome annotation pipeline for olive fly from this study and in comparison, to other available assemblies

Table S2: Assembly statistics of olive fly male and female chromosome individual chromosomes.

Table S3: Specific missing BUSCOs from the *Candidatus Erwinia dacicola* genome assembly. These missing BUSCOs are from the Bacteria and Enterobacterales conserved orthologue sets.

## Notes

### Competing Interest Statement

The authors have declared no competing interest.

